# Genomic potential and evolution of Dissimilatory Nitrate Reduction to Ammonium in Cyanobacteria

**DOI:** 10.1101/2023.05.23.542008

**Authors:** Manisha Ray, Shivakumara Manu, Gurdeep Rastogi, Govindhaswamy Umapathy

## Abstract

Cyanobacteria play an important role in primary production and nitrogen fixation. Although Cyanobacteria are well-known diazotrophic organisms, their role in other steps Nitrogen Cycle is obscure. Screening of Cyanobacterial genomes from cultured and unculturable species can help identify potentially novel functions. In this study, we assembled Cyanobacterial genomes from metagenomic data generated from environmental DNA isolated from a brackish water lagoon (Chilika, India). We annotated these Cyanobacterial metagenome-assembled genomes (MAG) for all the encoded functions using KEGG Orthology. We found two high-quality Cyanobacterial MAGs containing the *nirBD* gene and *nifH* and *nifD* genes involved in the nitrogen cycle. *nirBD* encodes for the Dissimilatory Nitrate Reduction to Ammonium (DNRA) activity, a function previously not ascribed to Cyanobacteria. We validated the presence of NirBD in publicly available isolate genomes of Cyanobacteria and examined its evolution in the phylum by phylogenetic reconciliation of species and gene trees. Our analysis revealed that both horizontal gene transfers and speciation events contributed to the dispersal of the *nirBD* gene in Cyanobacteria. We observed that mostly filamentous Cyanobacteria served as ancestral donors in horizontal gene transfer events. Further, we found that the *nirBD* gene is under a purifying selection pressure in Cyanobacteria. This study demonstrates the genomic potential and evolution of DNRA activity in Cyanobacteria for the utilisation of nitrate in the ecosystem which can help these organisms to cope with extreme environmental conditions. It expands our overall comprehension of the contribution of Cyanobacteria in the biogeochemical cycling in aquatic ecosystems.

## Introduction

Phylum Cyanobacteria account for 20-30% of Earth’s primary photosynthesis (Pisciotta et al., 2010) and are major primary producers in aquatic ecosystems. While some Cyanobacterial species are known for their ability to fix Nitrogen (N) but their involvement in other processes in the Nitrogen cycle is not well understood. One of these processes is dissimilatory nitrate reduction to ammonium (DNRA), in which nitrate serves as an electron donor and is converted to nitrite and ammonium (NO_3_ ^−^ → NO_2_ ^−^ → NH_4_ ^+^), providing a bioavailable N source to plants and animals (Lam & Kuypers, 2011). DNRA activity has been reported in Proteobacteria namely *Beggiatoa alba* (Vargas & Strohl, 1985) but its presence in Cyanobacteria is unclear. A recent study (Tee et al., 2020) indicated the expression of nirBD by *Microcoleus* in biofilms produced by them. DNRA has been reported to occur in both aerobic (Huang et. al 2020) and anaerobic conditions (Kamp et al., 2011). Further, it was also demonstrated through 15N isotopic studies that nirBD encodes for nitrate reductase and controls the DNRA process in Pseudomonas putida Y-9 (Huang et al., 2020).

Currently, more than 99% of bacteria have not been isolated in pure culture due to our limited understanding of the complex metabolic requirements necessary for their growth and cultivation in laboratory conditions (Yarza et al., 2014). Further, to understand the functional diversity of bacterial species in its entirety, we need to have whole-genome sequence information already available. But, to date, we have around 1.34 million bacterial species genomes in the NCBI database of which only 0.28% belong to Phylum Cyanobacteria whereas 72% of them belong to Phylum Proteobacteria. Of this, 0.28% (3900) of available Cyanobacterial genomes, 89% (3450) are from five of the total nine orders of Cyanobacteria. This shows that the genome sequence information in the present databases is skewed and Cyanobacterial species are severely underrepresented. This further exemplifies that only a few areas have been the focus of research like their ability to fix nitrogen and oxygenic photosynthesis. The sparse representation of some taxa in these genomic databases stems from the tedious and elaborate process of obtaining their axenic cultures and well as the physiology of many cyanobacterial species being symbiotic in nature (Waterbury, 2006). This “microbial dark matter” can be partially studied through the use of MAGs (Metagenome-Assembled Genomes) obtained from metagenomic data. In a study by Broman et. al. 2022 on a brackish water lagoon in the Baltic Sea, metagenomes were assembled and genes were predicted. They found that environmental variations played a more prominent role than the taxonomy of bacteria in the composition of functional genes. In another study, the evolutionary link between anaerobic basal thaumarchaeal lineages and mesophilic marine ammonia-oxidising archaea was found by reconstruction of two MAGs from phylum Thaumarchaeota (Reji & Francis, 2020).

Our study aimed to understand the evolution of DNRA function which has never been attributed to Cyanobacteria and is encoded by *nirBD* gene. We assembled Cyanobacteria genomes from environmental metagenomic data and annotated for hundreds of functions. Two of the high-quality MAGs showed the presence of the NirBD protein. Further, we deciphered the evolutionary history of DNRA function in Cyanobacteria by elucidating how this function was acquired and disseminated among species. Our study demonstrates the genomic potential and evolution of DNRA in Cyanobacteria for the utilisation of nitrate in the ecosystem.

## Methods

### 1) Study site, water sample collection, eDNA isolation and sequencing

Chilika Lagoon (19°28′ N: 19°54′ N & 85°06′ E: 85°35′ E) is a dynamic brackish coastal ecosystem with an average water spread area of 900 km2 and a catchment area of ∼4146 km2 (Srichandan et al., 2015). This lagoon is a Ramsar site and is a major biodiversity hotspot in India. Freshwater enters the Chilika Lagoon through 12 tributaries of Mahanadi river, while saline water intrudes into the lagoon from the Bay of Bengal, causing significant variability in physicochemical parameters due to the mixing of water of different masses (Tarafdar et al., 2021).

The sampling and sequencing of environmental DNA were carried out as previously described (Manu and Umapathy, 2023). Briefly, we filtered water from different locations of Chilika Lake, Odisha, India during the period from December 2019-December 2020 (Supplementary 1). For this purpose, we used a 0.45 µm mixed-cellulose ester filter membrane of 47mm diameter to trap all the extracellular DNA in the water so that the diversity of species and their functions are not overestimated by the ones which are abundant in cell numbers or their cell-lysis is easier. We used eDNA sampler by Smithroot Inc. (Thomas et al., 2018) and had 16 samples from 9 points from different seasons (Supplementary 1) in triplicates to maximise the yield. We used a lysis-free phosphate buffer-based eDNA isolation protocol from the filter membranes (Liang & Keeley, 2013) and Taberlet et al., 2012) We randomly fragmented the input DNA into 350 bp regions following which we used the Illumina Truseq DNA PCR-free library preparation method and sequenced for 300 cycles on the Novaseq 6000 platform.

### 2) Sequence analysis

#### 2.1) Assembly and taxonomic and functional annotation of MAGs

The raw reads were adapter trimmed and quality filtered with a phred quality threshold of 10 using the BBDUK tool (Bushnell, 2014). The quality-filtered reads were in-silico normalized to a target depth of 100x using BBNORM and error-corrected with BBCMS from the BBTOOLS package. The error-corrected reads from all samples were co-assembled using MEGAHIT with the meta-large pre-set. Contigs shorter than 1000 bp were filtered out and the quality-filtered reads from each sample were mapped to the remaining contigs with a minimum of 97% identity using BBMAP (Bushnell, 2014). The normalized abundance of contigs in all the samples was calculated using the JGI summarize bam contigs utility and the contigs were binned using METABAT2 (Kang et al., 2019) with default parameters. The quality of bins was assessed with CheckM (Parks et al., 2015) using the lineage workflow and bins with greater than 50% completion and less than 10% contamination were assigned as MAGs and retained for taxonomic classification. The MAGs were taxonomically annotated using the GTDB Toolkit (v. 207_2) with default parameters and those belonging to the Cyanobacteria were selected for functional annotation.

#### 2.2) Extraction of NirBD sequences from MAGs and curation of sequences of NirBD from the database

We generated the protein sequences from all the Cyanobacteria MAGs using Prodigal.v2.6.3 (Hyatt et al., 2010). All the predicted ORFs were annotated with KofamKOALA (Aramaki et al., 2020) which were then searched for DNRA using the definitions from the KeggDecoder tool. The definition for DNRA was *nirBD* (K00362 + K00363) and/or *nrfAH* (K03385 + K15876). Using this definition, we searched all the ORFs annotated by KofamKOALA. The e -value cut-off was set to more than 1e^-10^ and the score should be more than the threshold. These hits were searched using blastp to confirm their identity or to find the nearest match to the existing database. All the species having per cent identity greater than 50% and having both NirB and NirD proteins were selected for the construction of phylogeny. The protein family of the NirBD sequences was classified using InterPro (Blum et al., 2021). We considered all the database species of the protein family given that they had both *nirB* and *nirD* counterparts and their whole genomes are available in the database for constructing species trees using the ribosomal proteins.

##### a) Classification of proteins encoded by the MAGs into Cluster of Orthologous Groups (COGs)

To understand the functions encoded by the two MAGs that contain the NirBD protein, we utilized moshi4/COGclassifier (Shimoyama Y, 2022) to classify all the proteins into Cluster of Orthologous Groups (COGs). The Cluster of Orthologous groups categorizes the proteins encoded by the genome into specific functional categories, which can provide insight into the percentage of the genome that contributes to a particular functional class. Additionally, we manually searched for genes involved in the Nitrogen Cycle (from the Bacterial Kingdom based on available literature) among the hits predicted by the COGclassifier.

##### b) Construction of Species and NirBD protein Tree

###### Species Tree

The whole genomes of all the representative species were downloaded from NCBI (National Center for Biotechnology Information) database (Supplementary 2). We used GraftM (Boyd et al., 2018) to extract 15 single-copy ribosomal protein marker genes (Table 1) from all the representative species from the database as well as from the two MAGs generated as a part of this study. These 15 proteins were then aligned using MEGA11 (Tamura et al., 2021). These were then trimmed using Gblocks 0.91b (Castresana, 2000) and concatenated using MEGA11. A maximum-likelihood tree was constructed using an IQ-Tree webserver (Nguyen et al., 2015) using simultaneous model selection. For the construction of the Bayesian tree, PartitionFinder2 (Lanfear et al., 2017) was used to select the model for every partition followed by MrBayes 3.2.7a (Huelsenbeck & Ronquist, 2001).

**Table 1:**
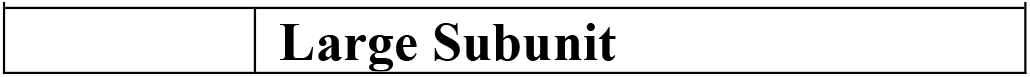
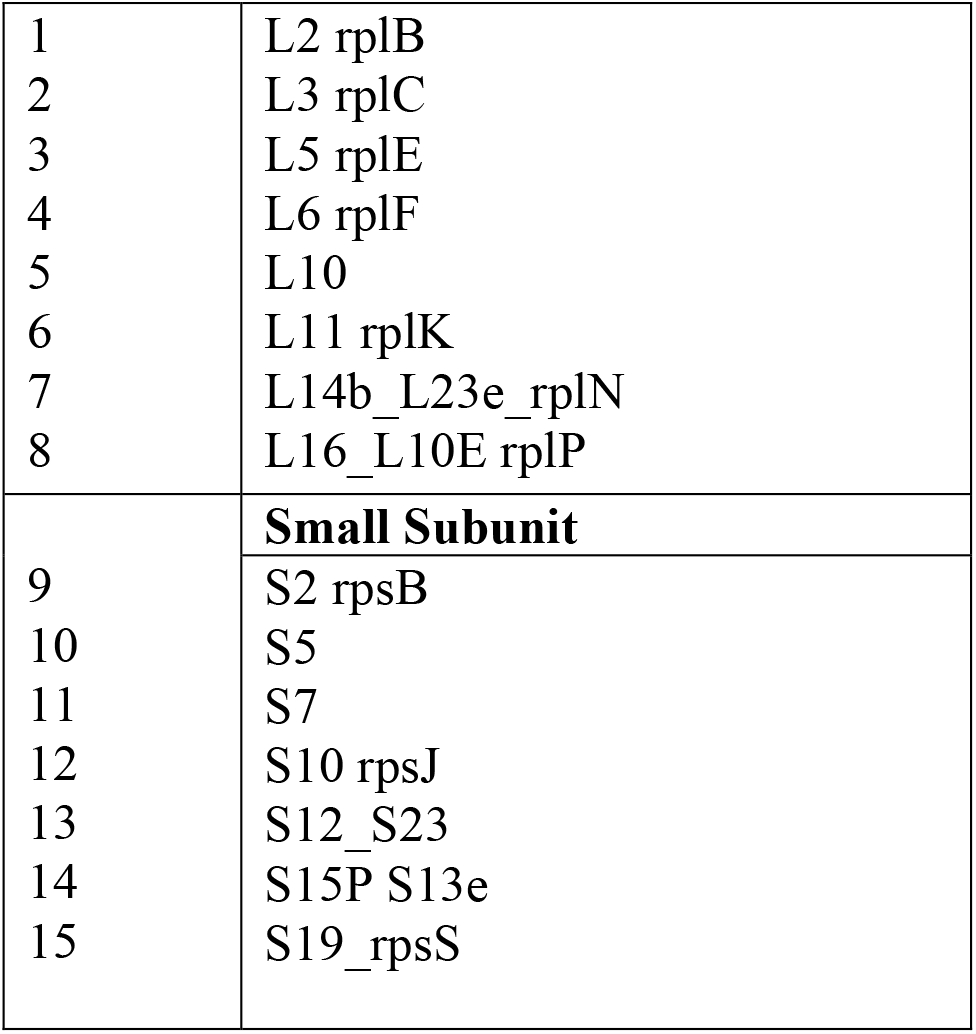
15 ribosomal proteins used in this study to construct Species Tree.

###### NirBD protein Tree

Blastp of the NirB and NirD ORFs from the two MAGs identified closest representative species having this protein in the database. These sequences were downloaded and aligned using MEGA11, followed by trimming using Gblocks 0.91b and concatenating using MEGA11. The same procedure was used to construct the maximum-likelihood and Bayesian tree as described for the species tree.

##### c) Reconciling of Species and NirBD protein tree

The topology of the phylogenetic tree for both species and the NirBD protein was inspected manually. Since there were several points of discordance between the two, a reconciliation was performed between species and NirBD protein trees by RangerDTL v2.0 (Bansal et al., 2018) to reconcile species and NirBD protein tree using default costs for duplication (2), transfer (3) and loss (1) for 200 reconciliations. These reconciliations were aggregated using RangerAggregator to obtain the most-supported events. Individual reconciliation was converted using RecPhyloXML converter (Python script) and visualised using Thinkind (Penel et al. 2022) webserver. The events that show more than 90% consistency of the event, as well as more than 90% mapping consistency, were depicted in the trees.

##### d) Selection analysis on NirBD protein

We used codeml (Yang, 2007), as implemented in EasyCodeML v1.4 (Gao et al., 2019) using Preset mode for the branch-model parameter. This model allows the ratio of non-synonymous (dN) to synonymous substitution rates (dS) called omega (ω) to vary among the branches of the phylogenetic tree to understand the impact of natural selection on this protein encoding for DNRA activity. The species tree in Newick format and aligned protein-coding DNA sequence of *nirBD* were used as input for EasyCodeML v1.4.

## Results

We assembled a total of 83 MAGs from Cyanobacteria of which 14 were high-quality and rest were medium-quality from Chilika Lagoon. We annotated the functions of these MAGs using KEGG Orthology and found that 7 of the MAGs had genes responsible for nitrogen-fixation. 2 of these 7 MAGs also showed presence of genes for DNRA activity as well.

### 1) Cyanobacterial genomes having NirBD signatures

The two Cyanobacteria MAGs showing the presence of the *nirBD* gene belonged to high-quality bins with less than 5% contamination > 90% completeness (Table 2).

**Table 2:**
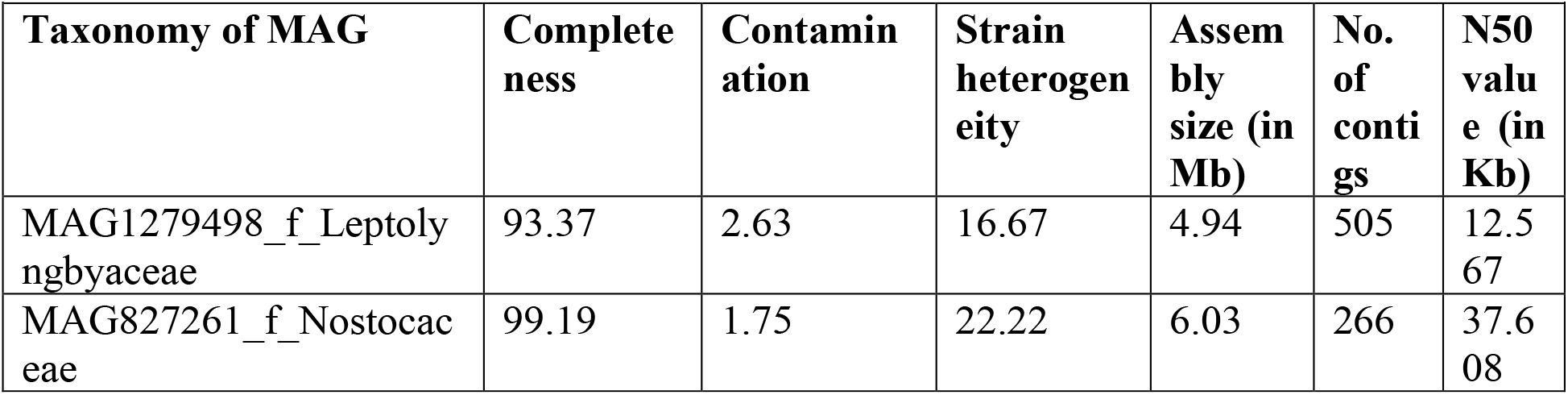
Detailed description of assembled genomes.

| Taxonomy of MAG | Completeness | Contamination | Strain heterogeneity | Assembly size (in Mb) | No. of contigs | N50 value (in Kb) |
| --- | --- | --- | --- | --- | --- | --- |
| MAG1279498_f_Leptolyngbyaceae | 93.37 | 2.63 | 16.67 | 4.94 | 505 | 12.567 |
| MAG827261_f_Nostocaceae | 99.19 | 1.75 | 22.22 | 6.03 | 266 | 37.608 |

The two MAGs showed the presence of a signature for DNRA activity-*nirBD* (K00362 + K00363) as depicted in Table 3. Thereafter upon classifying the obtained NirB and NirD protein sequences, they belonged to Nitrite reductase [NAD(P)H] large subunit (IPR017121) and Nitrite reductase (NADH) small subunit (IPR017881) protein families, respectively. We considered all the species belonging to these two protein families from Cyanobacteria. There were 38 species which had both IPR017121 and IPR017881 sequences available. We checked for genome sequences for all these Cyanobacterial species in the NCBI database and finally ended up with 32 genomes. Using the blastp search, we compared the NirB and NirD sequences of the MAGs and identified several Proteobacteria with a percentage identity of greater than 65%. The hits from Cyanobacteria in the blastp search were from the same genera as what we obtained from InterPro. We used these 43 sequences from Phylum Cyanobacteria and Proteobacteria and the two of our assembled MAGs for subsequent analysis.

**Table 3:** e-value scores for all the ORFs in the two MAGs upon scanning by KofamKOALA.

| Taxonomy of MAG (generated as part of this study) | KofamKOALA ID (KO) and query length | KO definition | e-value |
| --- | --- | --- | --- |
| MAG1279498_f_Leptolyngbyaceae | K00362<br>(835 amino acids) | Nitrite reductase<br>(NADH) large<br>subunit<br>[EC:1.7.1.15] | 0 |
|  | K00363<br>(110 amino acids) | Nitrite reductase<br>(NADH) small<br>subunit<br>[EC:1.7.1.15] | 2.7e-49 |
| MAG827261_f_Nostocaceae | K00362<br>(836 amino acids) | Nitrite reductase<br>(NADH) large<br>subunit<br>[EC:1.7.1.15] | 0 |
|  | K00363<br>(108 amino acids) | Nitrite reductase<br>(NADH) small<br>subunit<br>[EC:1.7.1.15] | 6.1e-45 |

### 2) Classification of Cluster of Orthologous Groups (COGs) in the two MAGs

#### a) MAG827261_f_Nostocaceae

moshi4/COGclassifier predicted a total of 5397 COGs for the MAG827261 belonging to the family Nostocaceae. Approximately 67% of the total COGs were attributed to specific functional classes and for the rest, only general function was predicted. Figure 1 (a) and (b) represent the proportion of COGs belonging to each functional category. Most of the COGs belonged to either signal transduction pathways (T) or cell wall and membrane biogenesis (M). We searched for the genes involved in the Nitrogen cycle among the predicted functional categories. We found the presence of NifD (Nitrogenase Mo-Fe protein) with an e-value of 1.03E-55 and NifH (Nitrogenase ATPase subunit) with an e-value of 2.55 E-133 in the COG category of Coenzyme transport and metabolism (H).

**Figure 1(a):**
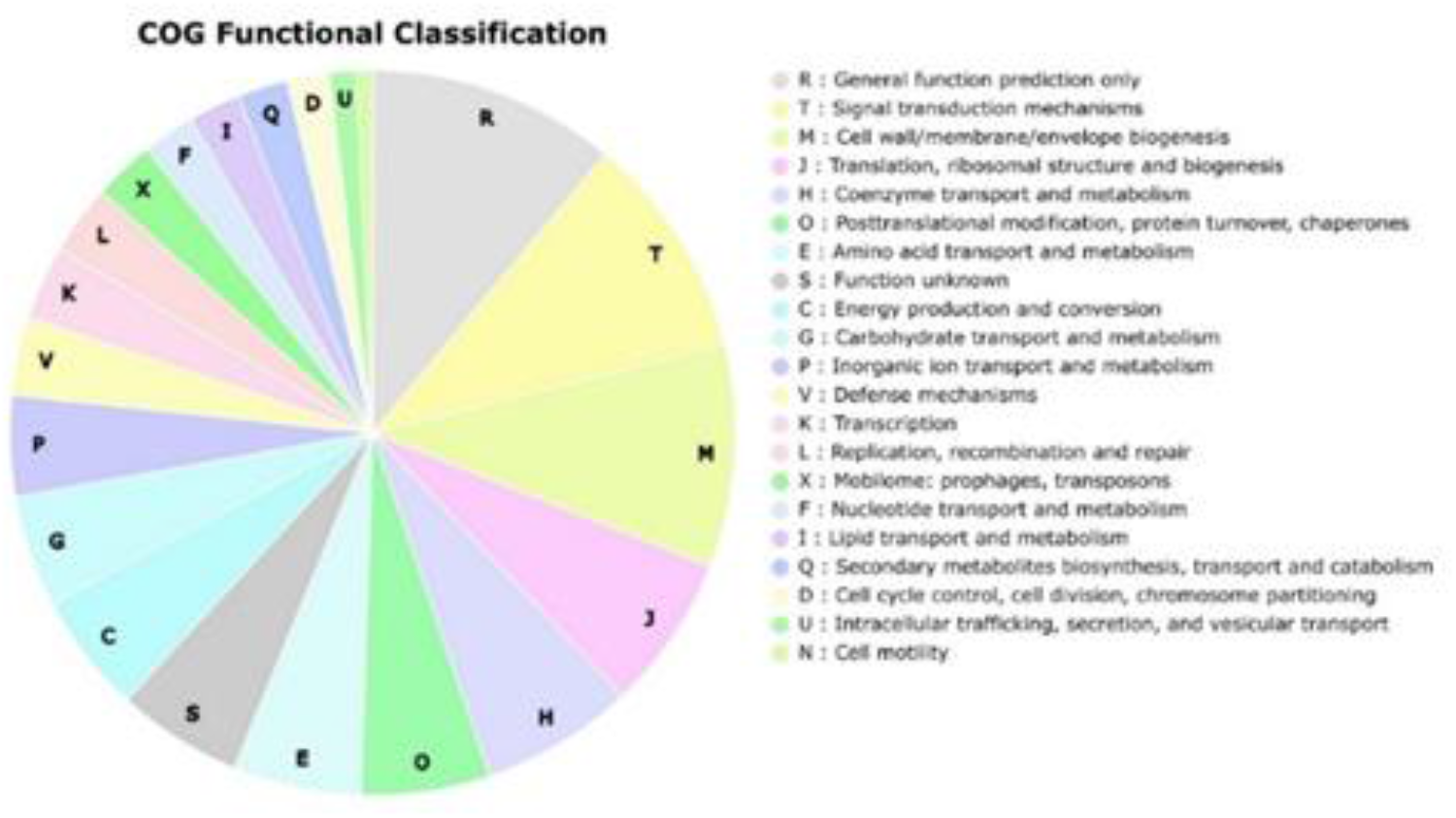
Pie-chart representation of the COGs encoded by MAG827261_f_Nostocaceae. The area in the pie-chart corresponds to the number of COGs belonging to that category.

**Figure 1(b):**
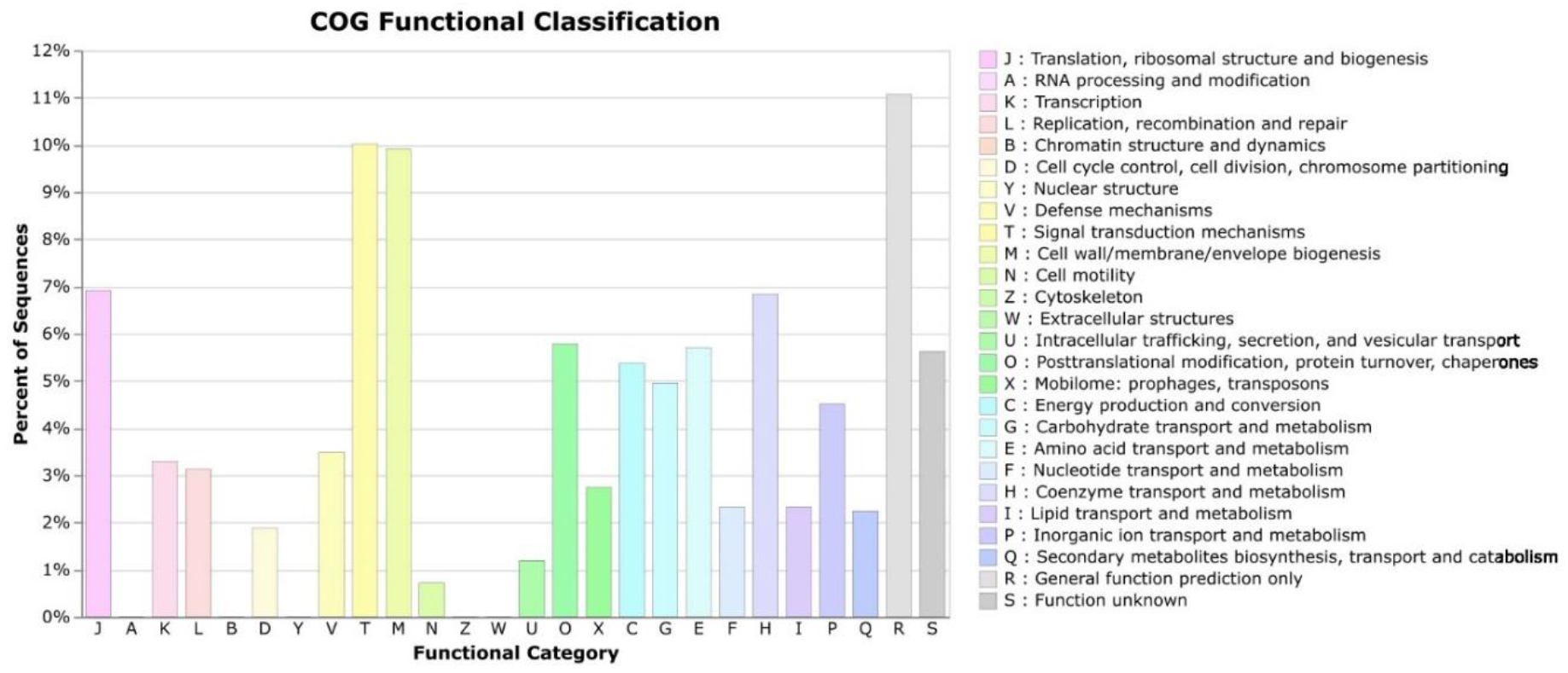
Bar-graph representation for the percent of COGs belonging to every category in MAG827261_f_Nostocaceae.

#### b) MAG1279498_f_Leptolyngbyaceae

4519 COGs were predicted by moshi4/COGclassifier for the MAG 1279498 belonging to the family Leptolyngbyaceae. Approximately 65% of the total of COGs were attributed to specific functional classes and for the rest, only general function was predicted. Figure 2 (a) and (b) represent the proportion of COGs belonging to each functional category. Most of the COGs belonged to either signal transduction pathways (T) or cell wall and membrane biogenesis (M). We searched for the genes involved in the Nitrogen cycle among the predicted functional categories. We found the presence of NifD (Nitrogenase Mo-Fe protein) with an e-value of 2.79E^-104^ and NifH (Nitrogenase ATPase subunit) with an e-value of 1.31 E^-178^ in the COG category of Coenzyme transport and metabolism (H).

**Figure 2(a):**
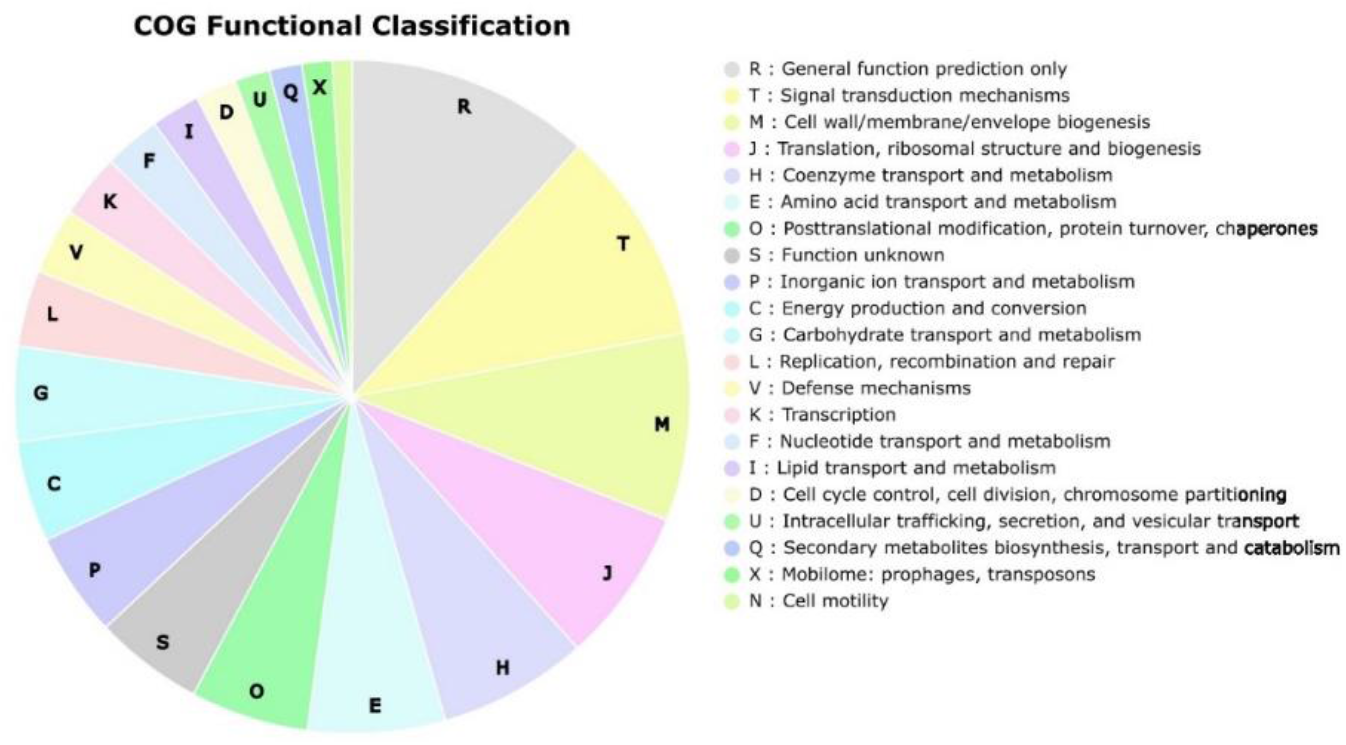
Pie-chart representation of the COGs encoded by MAG1279498_f_Leptolyngbyaceae. The area in the pie-chart corresponds to the number of COGs belonging to that category.

**Figure 2(b):**
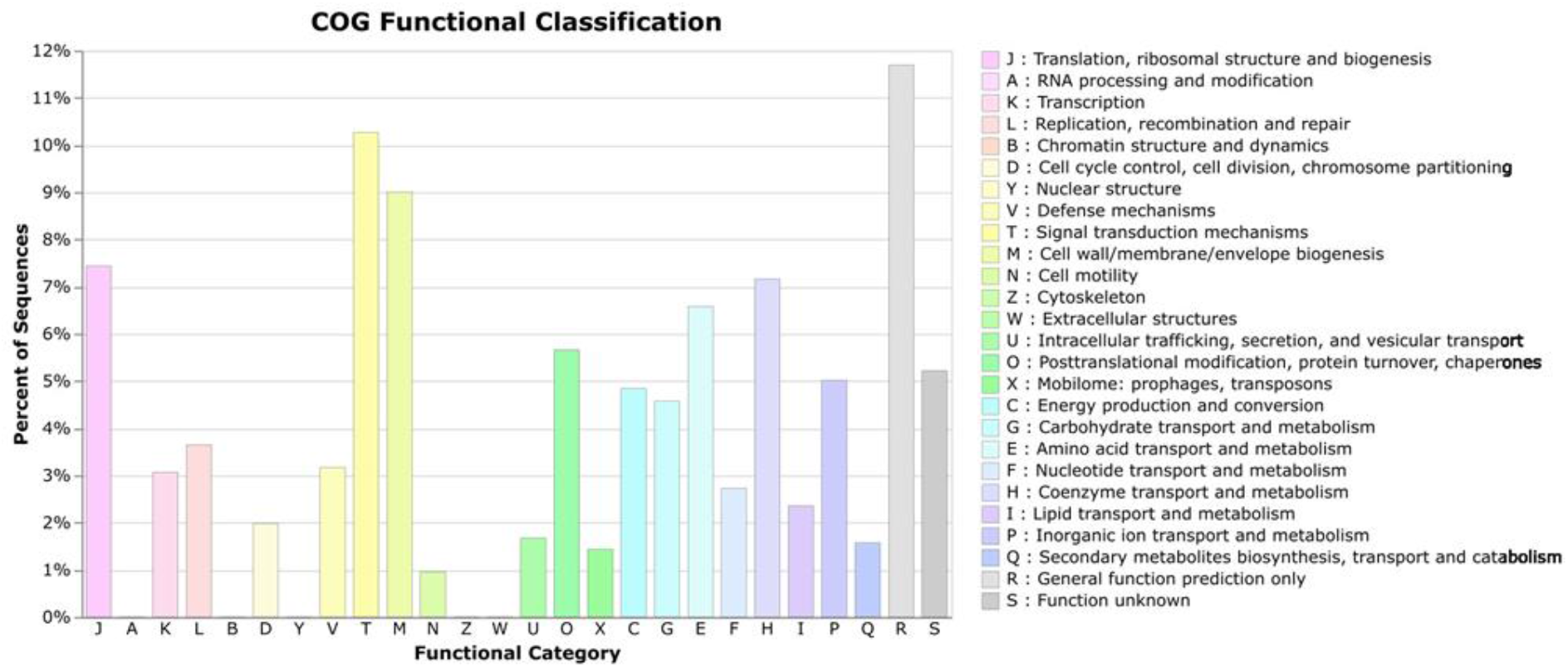
Bar-graph representation of the percent of COGs belonging to every category for MAG1279498_f_Leptolyngbyaceae.

### 3) Discordance between species and NirBD protein tree

We constructed the NirBD protein (Figure 3) and species trees (Figure 4) using both maximum likelihood and Bayesian methods. In both methods, the topology of the tree is the same but discordance is observed in several branches between the species and the NirBD protein tree. The species tree topology is consistent with the respective taxonomic assignments including the two assembled MAGs from Chilika Lake, India. But in contrast to this, we observe that NirBD protein phylogeny doesn’t follow the pattern as shown by organismal phylogeny. This signifies the evolutionary history of the NirBD protein did not happen concurrently with the evolution of the species. To decipher the reason and the source of these incongruencies, we performed a reconciliation analysis where we embedded the NirBD protein tree into the species tree. Further, this helps to deduce the gene losses, duplications and horizontal gene transfers which could be the probable reasons behind discordance.

**Figure 3:**
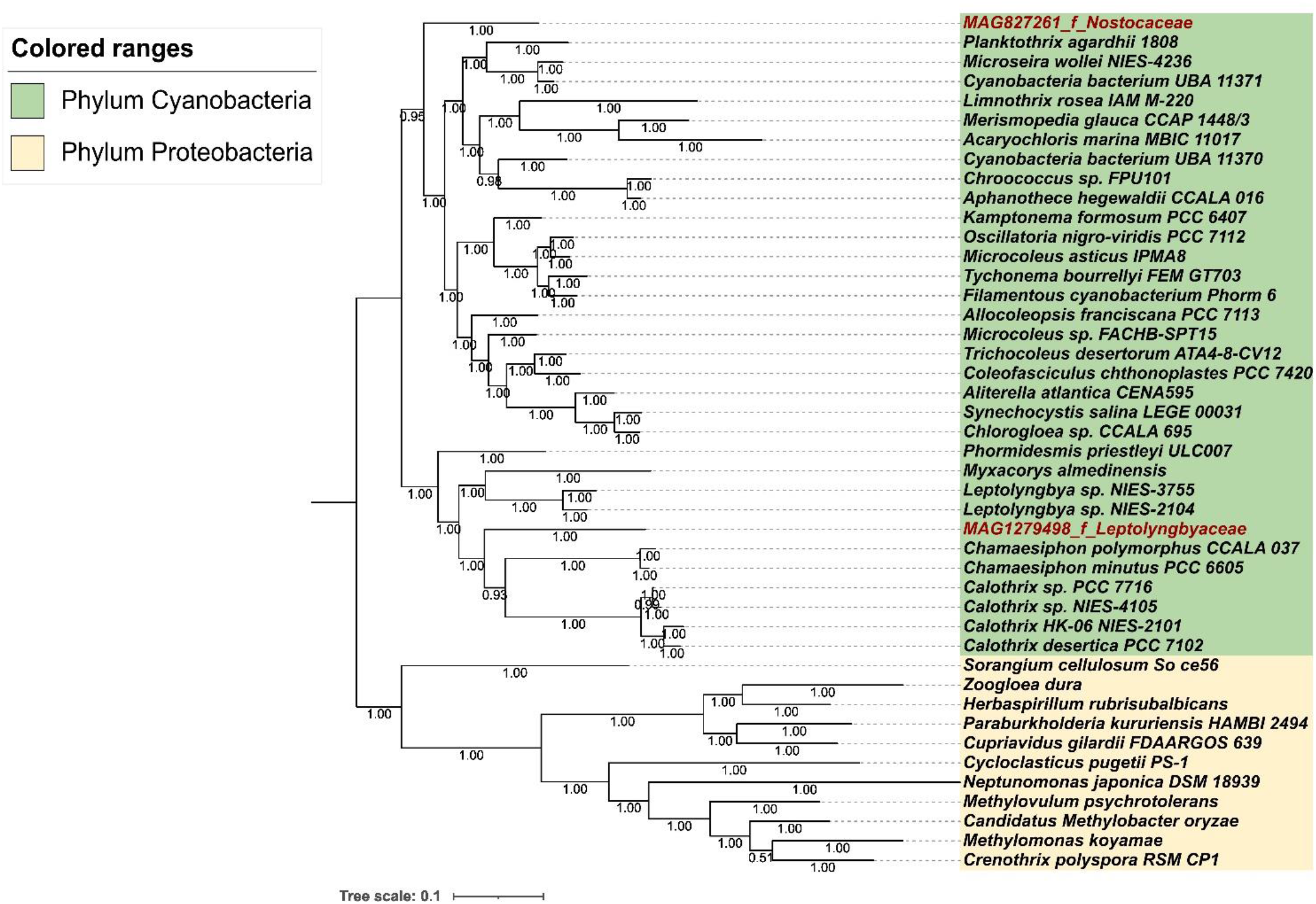
Concatenated NirB and NirD protein tree - The Bayesian Tree with bootstrap support values is depicted for each node. The NirBD extracted from MAGs constructed as a part of this study are represented in red text in this figure.

**Figure 4:**
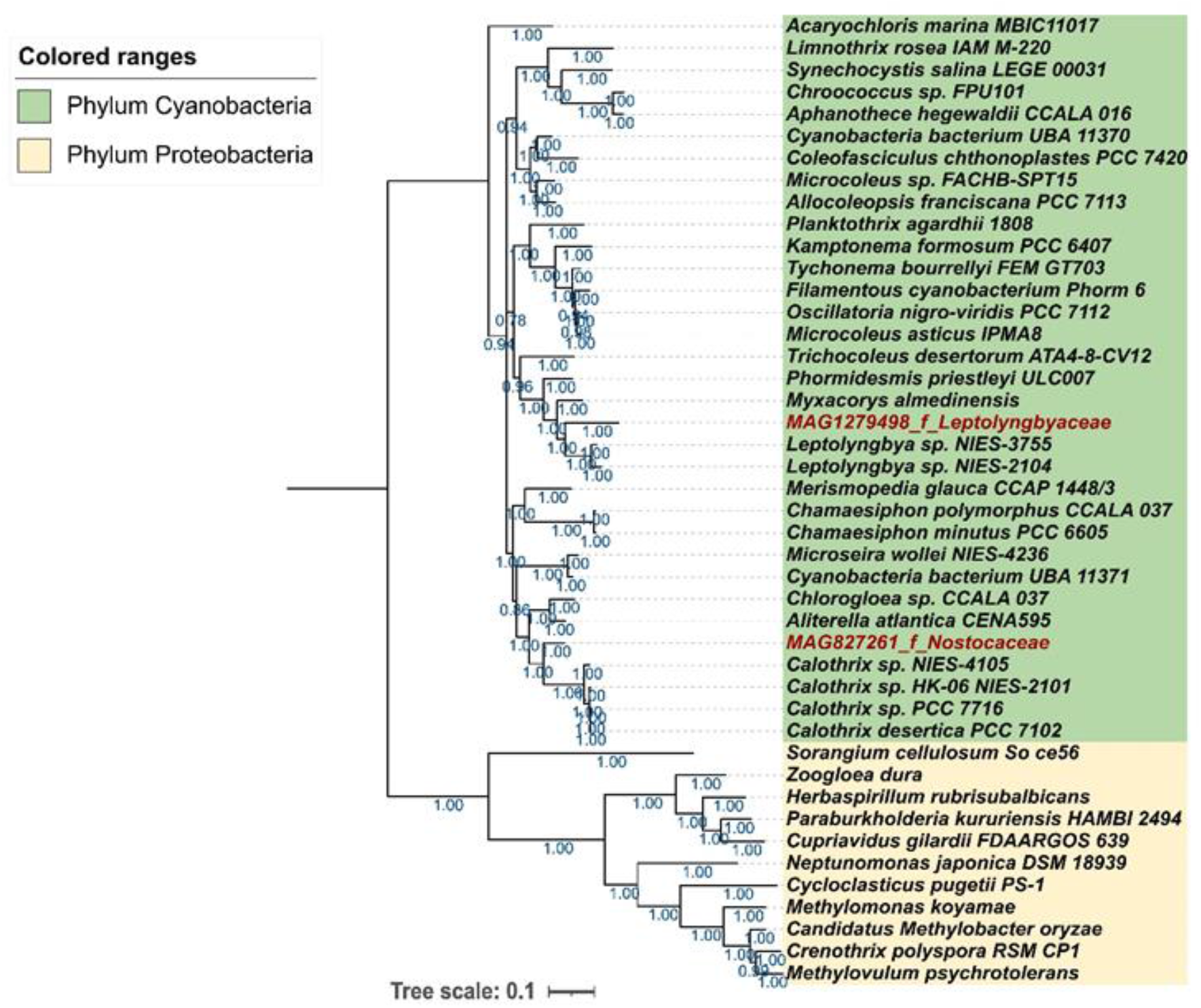
Species tree constructed using 15 concatenated ribosomal proteins and the Bayesian bootstrap values are mentioned for each node. The ribosomal proteins extracted from the MAGs generated as a part of this study are represented in red.

### 4) Reconciliation analysis

The dispersal of DNRA function traced by NirBD protein’s evolution demonstrates Cyanobacteria’s intra-phylum dynamics. This protein’s distribution results from a combination of horizontal gene transfer events and speciation among Cyanobacteria. Upon reconciling the NirBD protein tree with the species tree (Figure 5), we observe that the well-supported events with more than 90% consistency happened within the Cyanobacterial and not amongst their closest outgroup members from Proteobacteria. Most of the filamentous cyanobacteria have been responsible for the dissemination of the ability of DNRA. This is evident from the fact that the ancestral donor of NirBD (corresponding to node n22 in Fig. 3) has all of the filamentous species of cyanobacteria. Also, most of the HGT events are mapped to filamentous species of Cyanobacteria. Additionally, the deep-branching cyanobacterial species did not contribute to the dispersal of DNRA activity, implying that the event happened later during the evolution. The transfer events were mostly found to be restricted to specific species, indicating a limited ability to perform horizontal gene transfer. Additionally, it appears that most other species gained the NirBD protein through speciation from cyanobacterial ancestors that already possessed the potential for DNRA. The fact that the exact donor could not be identified in most cases suggests that the ancestral species with the potential for DNRA may not have their sequences represented in current databases, highlighting the limitations of an incomplete database.

**Figure 5:**
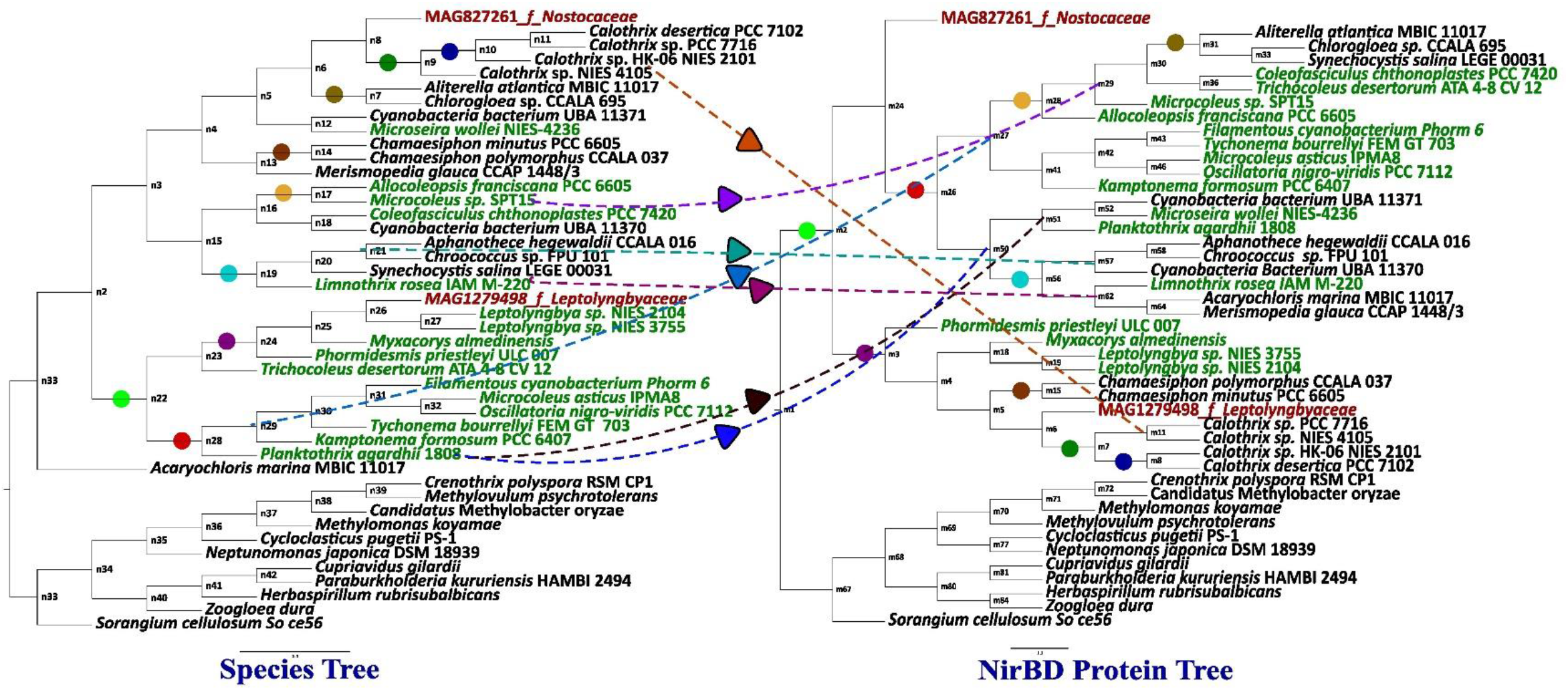
Reconciliation analysis of Species and NirBD protein tree. Only the events which were more than 90% consistent are shown here. The ones represented in green are filamentous cyanobacteria and the ones in red are MAGs constructed as part of this study. The different coloured circles illustrate the speciation event. For example, the red circle at n28 node of Species tree having six Cyanobacterial species speciated during the course of evolution and hence transferred the NirBD protein to ancestral species in m26 node of NirBD protein tree (marked in same colour). The horizontal gene transfer events are marked as dotted lines. Hence both the events could have been responsible for simultaneously disseminating this function.

### 5) Selection analysis

The ratio of dN/dS (ω value) signifies neutral selection if ω = 0, positive selection if ω > 1 and purifying selection if ω < 1. The LRT p-values and omega values for the selected branches are depicted in Table 5. The clade comprising 34 species of the Cyanobacteria shows a negative selection with an omega value of 0.0297 whereas the Proteobacterial outgroup comprising 11 species shows a positive selection with an omega value of 2.36188.

**Table 5:**
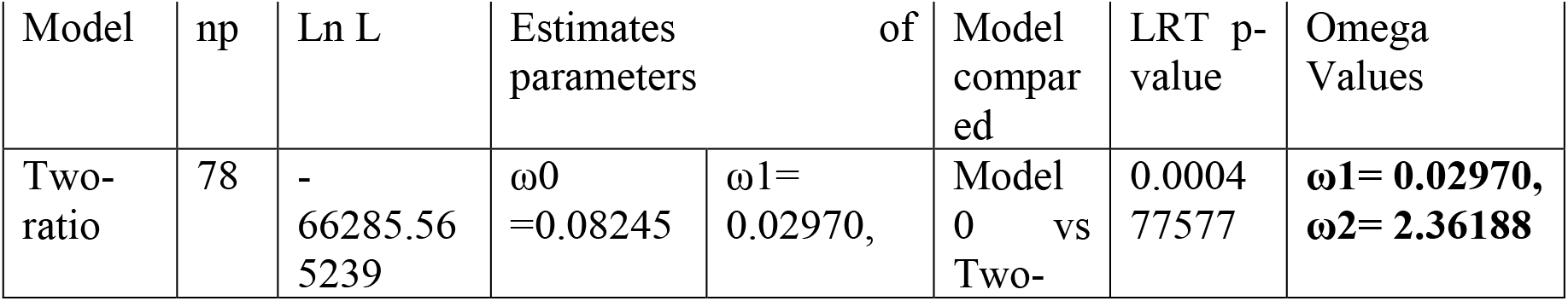
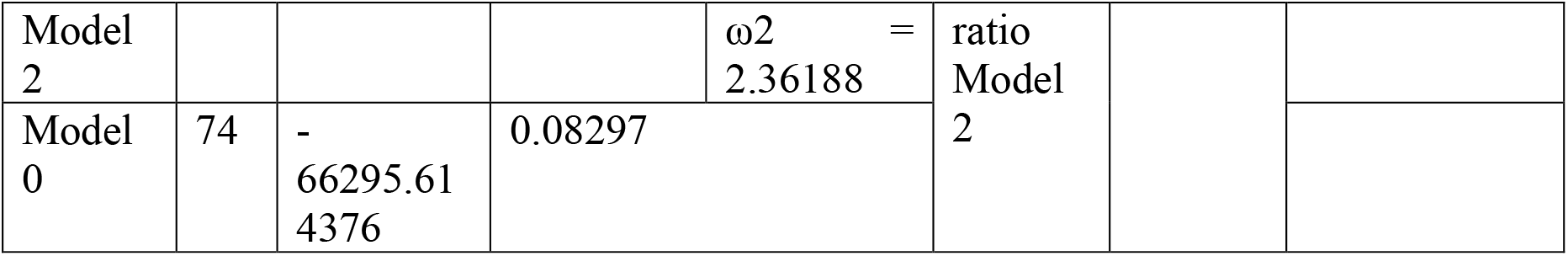
Branch Model (BM) results for codeml analysis.

## Discussion

DNRA is the conversion of nitrate to ammonium and its marker gene NirBD is reported in some bacterial species, but there is minimal information in Cyanobacteria many of which are diazotrophic. Hence, finding it in our assembled genomes from Chilika Lagoon was interesting but posed new questions. Upon subsequent analysis of Cyanobacterial whole-genomes from public databases (NCBI), and finding the NirBD gene in them suggested that NirBD gene in our cyanobacterial MAGs wasn’t any artefact and represents an unexplored area.

### Role of DNRA activity in Cyanobacteria

Some Cyanobacteria are well-known diazotrophs and are broadly classified into heterocystous and non-heterocystous forms. The enzyme nitrogenase, which is vital for nitrogen-fixing, is extremely sensitive to oxygen. Hence, heterocystous nitrogen-fixing Cyanobacteria has spatially parted the oxygenic photosynthesis from nitrogen-fixation by use of specialised cells called heterocyst. These specialised cells are devoid of oxygenic photosystem and possess a glycolipid cell wall aiding to minimise the oxygen concentration for nitrogen fixation to happen efficiently in these organisms (Stal and Zehr, 2008). In non-heterocystous nitrogen fixers, this is accomplished by temporal partitioning where nitrogen fixation happens during the dark when these species are grown under alternating light-dark cycles (Stal et al., 1994) and (Bergman et al. 1997). Interestingly, our analysis showed that many of the Cyanobacterial genera which are non-heterocystous nitrogen-fixers also had NirBD (a marker of DNRA function). Some of these genera were *Lyngbya, Phormidium, Oscillatoria, Microcoleus and Limnothrix*. We also aimed to look at all other functional COG categories and the genes involved in Biological Nitrogen Fixation (BNF) in our two Cyanobacteria MAGs. Both the MAGs showed the presence of NifD and NifH proteins and hence showing the ability to perform BNF besides DNRA. This hints towards the existence of multiple strategies in Cyanobacteria for acquiring bioavailable nitrogen sources (ammonium) and could be beneficial under certain stress conditions such as prolonged N-limitation. Our results of selection analysis pointed towards a negative selection of NirBD protein in Cyanobacteria. This could be because this function might be useful during stressed environmental conditions where other ways of synthesising bioavailable ammonia apart from DNRA are not possible. Under these conditions, having the potential to perform DNRA might serve as an advantage for the proliferation of the species.

### Discordance between Cyanobacterial species tree and NirBD protein tree

One of our assembled genomes (MAG827261_f_Nostocaceae), forms a separate clade in the NirBD protein tree demonstrating its amino acid sequence is quite divergent from the ones from the database. This MAG in the Species tree forms a sister clade with members of the *Calothrix* genus which also belongs to the family Nostocaceae (according to Genome Taxonomy Database, GTBD). This discordance in the phylogeny of the NirBD protein and the species (MAG827261_f_Nostocaceae) is due to their different evolutionary paths. Further, this also shows inadequate genome information in databases to fathom the evolution of these functional genes. This further leaves us with the resonant question about the indefinite diversity of such functions in ecosystems emanating from inadequate genome sequence information.

### Phylogenetic reconciliation and evolution of NirBD in Cyanobacteria

The genome of bacteria consists of core gene sets and flexible gene sets. The core gene sets are highly conserved in their amino acid sequence and are resistant to Horizontal Gene Transfers (HGT) and hence represent robust organismal phylogeny. Whereas flexible gene sets are variable and allow changes in their amino acid sequences according to selection pressure and can be transferred among organisms. In Cyanobacteria genomes genes encoding photosynthetic and ribosomal proteins form the densest component of core gene sets (Shi & Falkowski, 2008). Thus, the single-copy conserved ribosomal proteins are used to construct Species trees in bacteria since they are a reflection of the species evolutionary history (Hug et al., 2016) and (Moore et al., 2019). To understand the source of conflict between the phylogenies of a target gene tree and a Species tree, we need to comprehend their respective evolution. The discrepancies between a target gene tree with a species tree stem from the concept of independent evolution of species and the genes encoded by its genome. Embedding the gene tree into a species tree called phylogenetic reconciliation, can help in discerning the reason behind the discordance of the gene and organismal phylogeny (Waglechner et al., 2019) and (Shang et al., 2022). Analysing the phylogenetic reconciliation of our dataset illustrated that the deep-diverging nodes of the organismal phylogeny were not mapped as the donor for the NirBD protein. This meant that acquiring and disseminating NirBD protein happened later during evolution. Also, the *Acaryochloris marina* does not possess genes for nitrogen fixation as well (Pfreundt et al., 2012). Overall, we uncovered the genomic capacity and evolutionary dynamics of DNRA function within the Cyanobacteria Phylum.

### Future perspectives

Genome information serves as a blueprint of the range of functions that an organism has the potential to perform in different environmental scenarios. Meta-transcriptomics and proteomics can provide details of functions which are happening in real-time. This provides a clearer understanding of these functions and their importance in the ecosystem which changes biotic and abiotic interactions. Further, molecular assays to test the function of the target gene serve as a direct validation of it. Nonetheless, the integration of omics along with evolutionary studies can render thorough comprehension of these important functions.

## Conclusion

Our study revealed the genomic potential and evolution of DNRA function in Phylum Cyanobacteria. Some Cyanobacterial species possess alternate pathways for creating bioavailable ammonium in the ecosystem, which might confer a competitive advantage. While some Cyanobacterial species are known for Biological Nitrogen Fixation (BNF), research on other alternate pathways is limited. These studies open new avenues for the research community to further define the overall contribution of such functions in biogeochemical cycling and other important processes facilitated by microorganisms and when are they chosen by them. The rising interest in omics, particularly meta-transcriptomics and meta-proteomics, can help respond to these questions. This further reciprocates the roles of these microscopic creatures which go unnoticed but might be extremely helpful for ecosystem services.

## Data Availability

The raw reads from the metagenomic sequencing are available in NCBI SRA under the BioProject accession PRJNA691704.

## Acknowledgments

We acknowledge the support received by G.U. from Department of Biotechnology (DBT), Govt. of India, vide Grant No. BT/PR29032/FCB/125/4/2018 for this study which also included funding of PhD fellowship for M.R. S.M. was supported by a BINC fellowship from DBT, Govt. of India.

We thank Chilika Development Authority, Balugaon, Odisha for the help received during sample collection. We also thank Dr. Mihir Trivedi for his constructive suggestions during phylogenetic analysis.

## Supplementary material

Supplementary 1: Details of locations and period of sampling

Supplementary 2: Genomes of Cyanobacteria used in this study to construct Species Tree and NirBD protein tree phylogeny

